# Edge Community Entropy is a Novel Neural Correlate of Aging and Moderator of Fluid Cognition

**DOI:** 10.1101/2023.10.11.561957

**Authors:** Anita Shankar, Jacob Tanner, Tianrui Mao, Richard Betzel, Ruchika Shaurya Prakash

## Abstract

Decreased neuronal specificity of the brain in response to cognitive demands (i.e., neural dedifferentiation) has been implicated in age-related cognitive decline. Investigations into functional connectivity analogues of these processes have focused primarily on measuring segregation of nonoverlapping networks at rest. Here, we used an edge-centric network approach to derive entropy, a measure of nodal specialization, from spatially overlapping communities during cognitive task fMRI. Using Human Connectome Project Lifespan data (713 participants, 36-100 years old), we characterized a pattern of nodal despecialization differentially affecting the medial temporal lobe and limbic, visual, and subcortical systems. Global entropy uniquely covaried with age when controlling for network segregation. Importantly, relationships between both metrics and fluid cognition were age-dependent, although entropy’s relationship with cognition was specific to older adults. These results suggest entropy is a potentially important metric for examining how neurological processes in aging affect functional specialization at the nodal, network, and whole-brain level.

---

Like other complex systems^1^, the human brain exhibits hierarchical and multiscale organization that maximizes efficiency of information transfer while maintaining flexibility to changing demands and resiliency to damage^2^. This hierarchical organization also facilitates complex human behavior, including higher-level cognitive functions that show declines across the adult lifespan, such as attention, processing speed, and episodic memory^3–5^. These declines can lead to decreased quality of life, increased public health burden, and are considered risk factors for progression to pathological diseases such as Alzheimer’s disease and other dementias^6,7^. As such, there is growing interest in the field of clinical neuroscience to characterize network hierarchical organization and brain dynamics across the lifespan to explain and predict cognitive outcomes associated with aging and disease^8,9^.

One property of brain networks that demonstrates significant age-related changes is modularity, which refers to the degree to which the network is composed of sub-networks with strong intra-network connectivity and weaker inter-network connectivity^10–13^. Studies of resting-state functional magnetic resonance imaging (fMRI) have consistently shown age-related decreases in network segregation, including whole-brain modularity^14–17^, which is estimated by algorithmically partitioning the network into clusters or “modules” by maximizing the within-module density of connections compared to a null connectivity model^12^. These age-related changes at the whole-brain level are in part driven by reductions in within-network connectivity and increases in between-network connectivity and involve sub-networks such as the default mode network (DMN), somatosensory, and control networks^17–20^. Interestingly, these network changes have also been observed in Alzheimer’s disease and are associated with pathological changes in the disease^21–24^. Taken together, these results reflect a consistent pattern of neural desegregation throughout aging that has been suggested as a connectivity-based analogue of neural dedifferentiation^25,26^, a term that initially referred to age-related efficiency and fidelity of neuronal firing during processes supporting cognition^27–29^. More recently, neural dedifferentiation has been used more broadly to describe the brain’s de-specialization of function with increasing age and disease processes, especially where this decreased specialization is indicative of decreased cognitive performance^25,27^.

However, most studies describing age-related differences in network properties have focused on resting state data and non-overlapping definitions of brain modules, in which nodes are assigned to a single module only. Relatively few studies have characterized the brain’s overlapping modular structure, let alone described its variation across the lifespan^30–32^. Furthermore, few studies to our knowledge, have investigated the concept of functional specialization, rather than segregation, as a potentially important metric in aging and neural dedifferentiation. Here, we operationalize age-related neural dedifferentiation using task fMRI and a recently proposed edge-centric framework^33,34^. In this framework, time-invariant correlations (functional connections) are decomposed into their framewise contributions, yielding time-resolved (dynamic) estimates of connections’ weights. Based on their temporal similarity to one another, edges can be grouped into non-overlapping clusters, whose assignments can be projected back to individual nodes, revealing pervasively overlapping cluster structure. From the probability distributions of nodal participation in each of these clusters, or communities, we are able to calculate entropy— a measure used in information science^35^ indicative of the diversity of edge-community participation^33^ as a neuromarker of decreased specialization across the lifespan at the level of the individual node, network, and the whole brain.

Only recently have researchers started to investigate how entropy may have significance as a network communication measure in neuroscience, establishing that entropy varies across functional systems in ways that are distinct from other measures of modular structure, that entropy can be modulated by tasks such as movie watching, and that entropy changes over the stroke recovery process^33,36^. Here, we hypothesize that as the brain ages and decreases in neural specificity result in less distinct functional neuronal populations and decreased spatial segregation of brain regions into modules^25,27^, these regions will also demonstrate increased diversity in community participation, as represented by their entropy.

Typically, studies linking network properties to cognition have examined whether age-related changes are indicative of neural compensation or dedifferentiation processes by correlating these metrics with cognition^28^. Here, we instead build upon recent work suggesting that some whole-brain network metrics, such as modularity, can also reflect network integrity, greater capacity for plasticity, and higher resilience to damage ^37–39^. Thus, our secondary aim in this study was to investigate global whole-brain entropy as a moderator of the cross-sectional relationship between increased age and decreased cognitive performance across the lifespan. We hypothesized that factors consistent with lower neural dedifferentiation (i.e., greater nodal specialization as measured by lower entropy) will be indicative of a network that is resilient to age-related changes and therefore demonstrates better cognitive performance. Furthermore, given the novelty of this metric in the aging literature, we compare entropy’s age and cognition-related variance to both modularity and participation coefficient, a conceptually similar nodal graph metric. To perform these analyses, we used fMRI data collected during three cognitive tasks (attention, visual-motor processing, episodic memory) and fluid cognition scores from the National Institutes of Health Cognitive Toolbox obtained from the Human Connectome Project Lifespan Release ^40^.

## Results

The primary aim of this study is to 1) investigate lifespan changes in entropy as a measure of nodal specialization during the performance of cognitive tasks, and 2) examine whether entropy moderates cross-sectional lifespan decreases in fluid cognition. To achieve these aims, we analyzed data from the Lifespan Human Connectome Project Aging (HCP-A) 2.0 Release. Our final dataset consisted of 713 adults, 55.7% of whom were female, ranging in age from 36 to over 100 years old (Table 1).

**Table 1.** Demographic, Cognitive, and Brain Information.

| Table 1 \| Demographic, Cognitive, and Brain Information |  |  |  |  |
| --- | --- | --- | --- | --- |
| Variables | Full Sample<br>M (SD) | Ages 36-64<br>M (SD) | Ages 65-79<br>M (SD) | Ages 80-100+<br>M (SD) |
| **Demographic** |  |  |  |  |
| Age | 60.36 (15.73) | 49.39 (7.97) | 71.54 (4.56) | 86.02 (5.93) |
| Sex |  |  |  |  |
| Female, n, % | 406, 56% | 250, 57.60% | 99, 52.94% | 57, 54.81% |
| Male, n, % | 319, 44% | 184, 42.40% | 88, 47.06% | 47, 45.19% |
| Race |  |  |  |  |
| White n, % | 525, 72.41% | 267, 61.52% | 160, 85.56% | 98, 94.23% |
| Black n, % | 101, 13.93% | 85, 19.59% | 12, 6.42% | 4, 3.85% |
| Asian n, % | 52, 7.17% | 39, 9.00% | 13, 6.95% | 0, 0% |
| More than one race n, % | 31, 4.28% | 28, 6.50% | 2, 0.01% | 1, 0.96% |
| Unreported n, % | 14, 1.93% | 14, 3.23% | 0, 0% | 0, 0% |
| Ethnicity |  |  |  |  |
| Hispanic or Latino n, % | 74, 10.21% | 66, 15.21% | 8, 4.28% | 0, 0% |
| Not Hispanic or Latino n, % | 649, 89.52% | 368, 84.79% | 178, 95.19% | 103, 99.04% |
| Unreported n, % | 2, 0.28% | 0, 0% | 1, 0.53% | 1, 0.96% |
| Education, years | 17.47 (2.22) | 17.46 (2.15) | 17.82 (2.13) | 16.97 (2.55) |
| **Cognitive** |  |  |  |  |
| NIH Toolbox Fluid Cognition | 99.11 (12.58) | 103.45 (11.47) | 94.82 (9.62) | 83.80 (10.41) |
| **Graph Theory Metrics** |  |  |  |  |
| Whole-brain task-averaged Entropy | 0.78 (0.03) | 0.77 (0.03) | 0.79 (0.03) | 0.81 (0.02) |
| Whole-brain task-averaged Part. Coefficient | 0.42 (0.06) | 0.41 (0.06) | 0.43 (0.05) | 0.46 (0.05) |
| Whole-brain task-averaged Modularity | 0.48 (0.04) | 0.51 (0.03) | 0.48 (0.04) | 0.44 (0.05) |
| **Other** |  |  |  |  |
| Task-averaged mean FD | 0.16 (0.08) | 0.15 (0.08) | 0.18 (0.09) | 0.20 (0.08) |
| Task-averaged mean FD<sup>2</sup> | 0.02 (0.03) | 0.02 (0.02) | 0.02 (0.03) | 0.03 (0.03) |
| Estimated total intracranial volume, mm<sup>3</sup> | 1.53e+06<br>(1.91e+05) | 1.49e+06<br>(1.92e+05) | 1.57e+06 (1.80e+05) | 1.59e+06<br>(1.73e+05) |
| Note: Descriptive statistics for demographic variables, cognitive performance, graph theory metrics, and covariates. Mean FD refers to the motion metric average framewise displacement. |  |  |  |  |

Throughout this section, we focus on findings that result from averaging graph theory metrics across individual cognitive tasks and include information about task-based differences as an exploratory analysis. Furthermore, although our focus is on the entropy measure and edge-centric analysis, we compare entropy to the more familiar network metrics of modularity and participation coefficient, contextualizing our findings with respect to the extant literature. Finally, our results refer to our findings for the Glasser atlas, developed specifically for the HCP dataset, unless otherwise noted. We replicated these analyses using a second functional atlas, the Schaefer 300 parcellation atlas, and reference results from this atlas when noteworthy.

### Entropy values vary across nodes, networks, and age groups

We calculated normalized entropy for each node, examining patterns of entropy across networks and the entire brain for the three HCP-Aging defined age groups: mature adults (ages 36-64), old adults (ages 65-79), and oldest-old adults (ages 80 and above). At the nodal level, we observed high and low entropy values dispersed across the brain. The lowest values were across the temporal lobe and insular cortex while the highest values were across areas of the parietal and occipital lobes in all three age groups (Figure 1a). By network, the median entropy value across age ranges was highest in visual and attentional networks and lowest in subcortical, ventral-multimodal, and orbito-affective systems (Figure 1b, 1c). For the Schaefer atlas, this corresponded to highest entropy in the visual central, dorsal attention, and control A networks, and lowest entropy in the limbic and subcortical systems (Supplementary Figure 1c). To determine whether high entropy occurs only in the context of old age, we examined the range of raw entropy scores across the lifespan. We found that even in the youngest group of adults (148 individuals aged 35-45), nodal entropy values ranged from 0.1003 to 0.9903, spanning close to the entire theoretical range (0-1).

**Figure 1.**
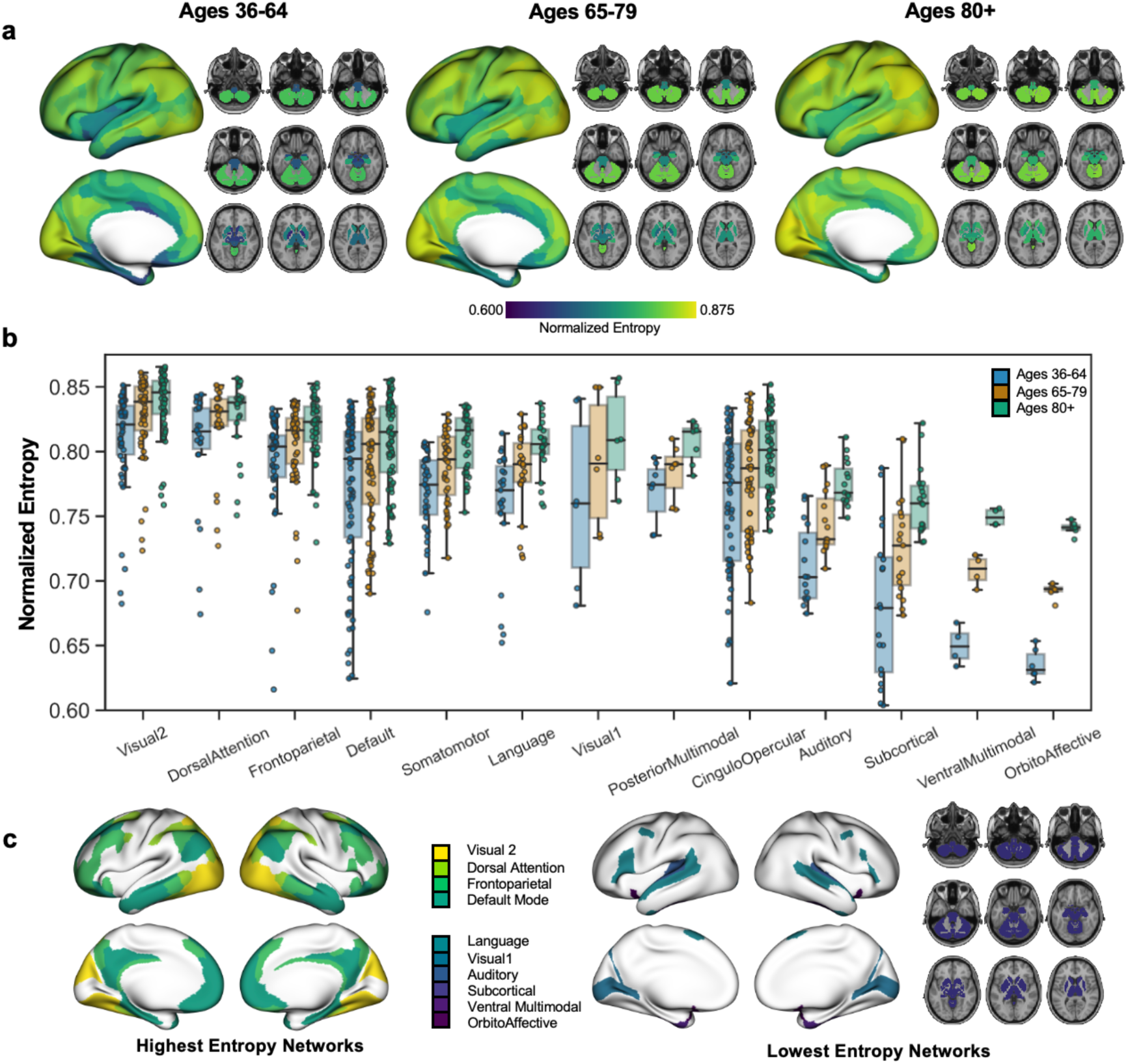
Normalized Entropy. **a.** Normalized entropy at the nodal level across the whole brain, averaged across HCPA-defined age groups of 36-64, 65-79, and 80-100+. **b.** Normalized entropy at the network level by HCPA-defined age groups, **c.** Networks with the highest normalized entropy values across the entire age range (Visual 2, Dorsal Attention, Frontoparietal, Default Mode), **d.** Networks with the lowest normalized entropy values across the entire age range (Language, Visual 1, Auditory, Subcortical, Ventral Multimodal, OrbitoAffective).

### Entropy increases with age and differentially presents in medial-temporal and subcortical brain regions and affects connectivity in multiple networks

In the present study, we were primarily interested in the relationship between entropy and age. To statistically test this relationship, we computed the bivariate, product-moment correlation between task-averaged entropy and age at the nodal, network, and whole-brain level. Consistent with our hypothesis that brain regions would, on average, demonstrate involvement in a greater proportion of communities with increasing age, we found a positive correlation between age and whole-brain entropy R(713)=0.53 *p*=6.02e-54 (Figure 2a). We also investigated this relationship controlling for neural covariates such as in-scanner motion and total brain volume (B = 0.46, *p* = 9.98e-41), and additionally for demographic factors including sex, race, ethnicity (B = 0.48, *p* = 8.36e-42) (Table 2a). These findings were similar for the Schaefer atlas (Supplementary Table 4).

**Figure 2.**
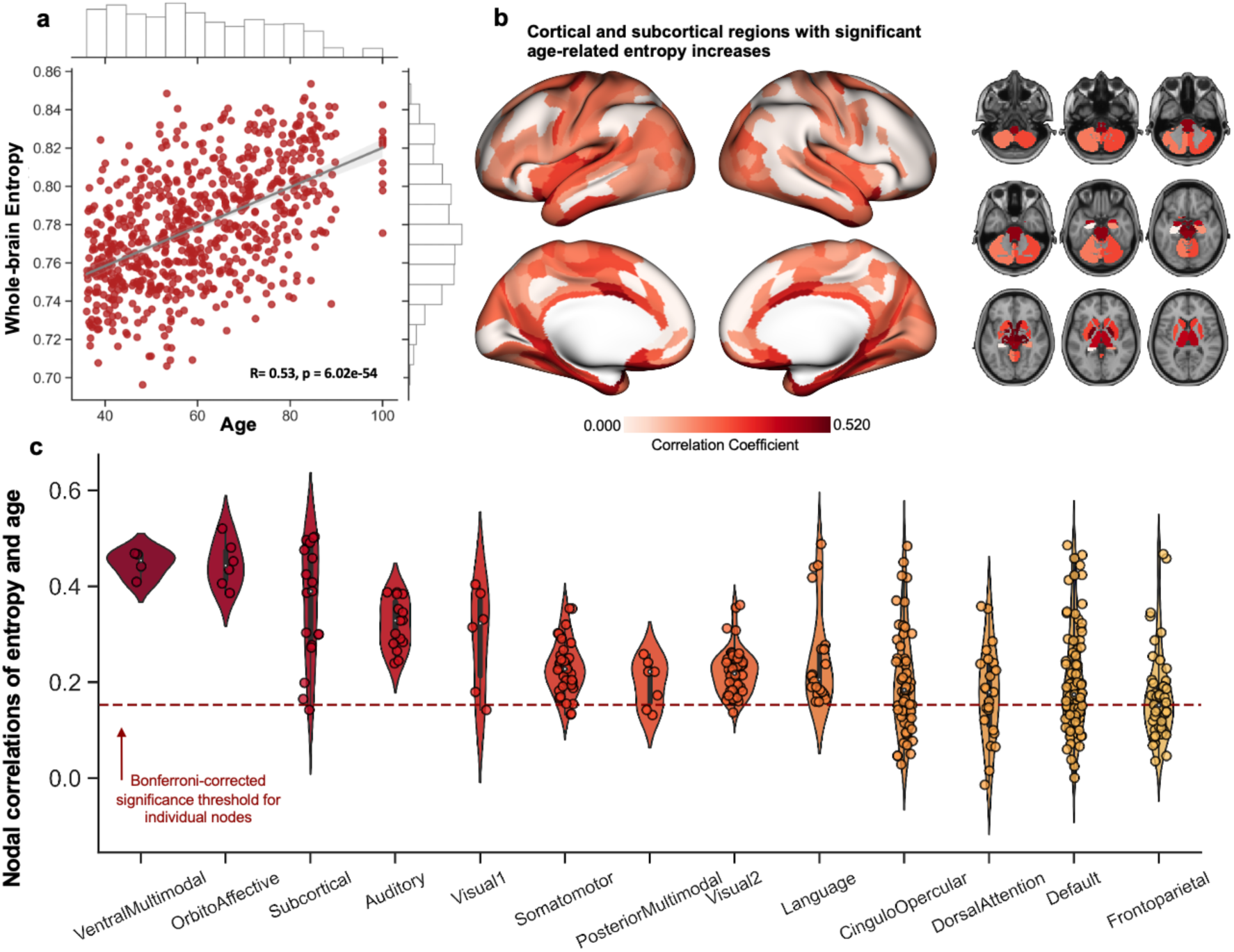
Entropy correlated with age. **a.** Scatter plot of correlation between averaged whole-brain entropy and age. **b.** Brain plots of nodes with significant age­entropy correlations, **c.** Violin plot of nodal correlations between age and entropy organized by network. Red dotted line indicates Bonferroni-corrected significance threshold for nodal relationships.

**Table 2a.**
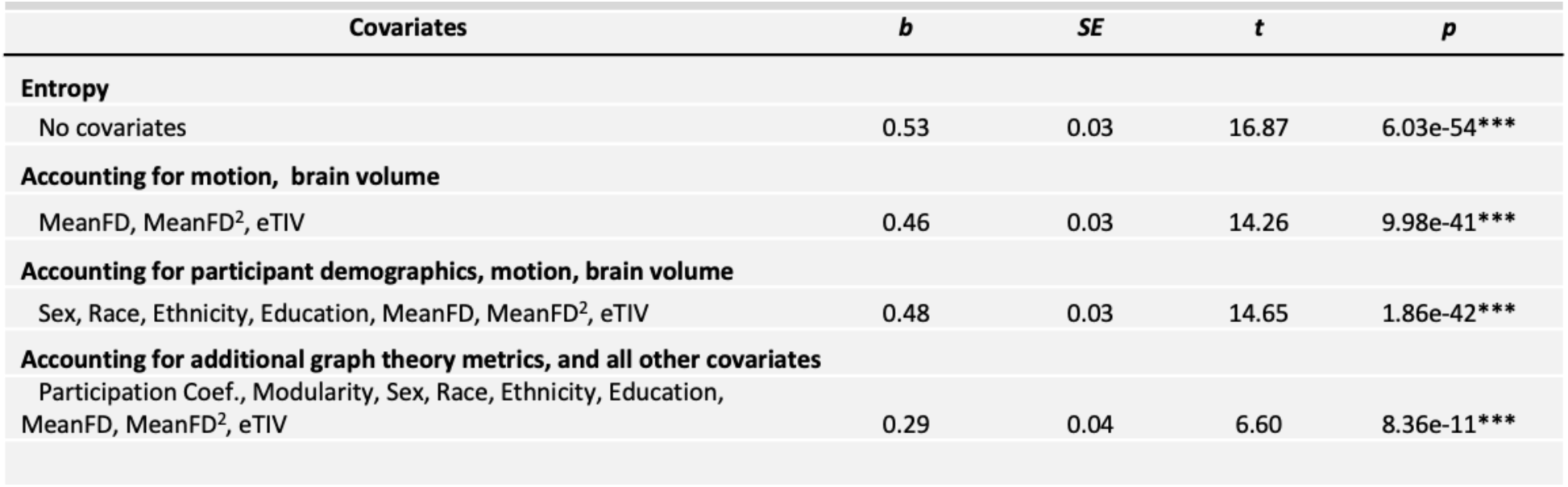
Linear regression statistics of relationship between whole-brain task-averaged entropy and age, accounting for covariates.

| Covariates | *b* | *SE* | *t* | *p* |
| --- | --- | --- | --- | --- |
| **Entropy** |  |  |  |  |
| No covariates | 0.53 | 0.03 | 16.87 | 6.03e-54*** |
| **Accounting for motion, brain volume** |  |  |  |  |
| MeanFD, MeanFD<sup>2</sup>, eTIV | 0.46 | 0.03 | 14.26 | 9.98e-41*** |
| **Accounting for participant demographics, motion, brain volume** |  |  |  |  |
| Sex, Race, Ethnicity, Education, MeanFD, MeanFD<sup>2</sup>, eTIV | 0.48 | 0.03 | 14.65 | 1.86e-42*** |
| **Accounting for additional graph theory metrics, and all other covariates** |  |  |  |  |
| Participation Coef., Modularity, Sex, Race, Ethnicity, Education, MeanFD, MeanFD<sup>2</sup>, eTIV | 0.29 | 0.04 | 6.60 | 8.36e-11*** |

We further examined regional differences in the relationship between entropy and age. We found that entropy significantly increased with age in 304 nodes (80.21%), with *p*-values Bonferroni corrected for 379 comparisons (Figure 2b, 2c). Nodes with the strongest age-related increases were found across several networks, including the default-mode, frontoparietal, cingulo-opercular, language, and orbito-affective networks. These results demonstrated that entropy does not significantly increase with age in all regions of the brain. Results further showed that even within significant increases, there is variability in the degree of increase, with significant correlations ranging from 0.15 to 0.52. We also investigated network-level age entropy. We found that all networks contained nodes with significant age-related changes to entropy. For the Glasser atlas, entropy in at least 50% of the nodes in every network significantly increased with age (Figure 2c). However, for the Schaefer atlas, this was true of 11 out of the 18 networks, with the greatest proportion of significant nodes in the limbic networks, subcortical regions, and parts of the default-mode network (Supplementary Figure 2c). Importantly, no networks and no individual nodes demonstrated a significant negative correlation with age, with a minimum nodal correlation of R = −0.01, *p* = 0.71, and even networks with high normalized entropy demonstrated significant entropy increases.

To examine the most consistent regions with entropy increases, we identified nodes with significant age increases in all three of the individual cognitive tasks (Supplementary Figure 6). We found 86 nodes (22.69%) in which entropy significantly increased with age across all three cognitive tasks, with *p*-values Bonferroni corrected for 1137 comparisons (379 nodes, 3 tasks) (Supplementary Figure 6d). These nodes appeared to be spatially adjacent to one another and consisted of 36 nodes (41.9%) in the temporal lobe, including the entire bilateral medial temporal lobe and auditory cortices, 21 nodes (24.4%) in the frontal lobe, primarily the anterior cingulate and medial prefrontal cortex, 21 nodes (24.4%) in the insula and subcortex, 4 (4.7%) nodes in the occipital lobe, and 4 (4.7%) nodes in the parietal lobe.

### Entropy demonstrates unique age-related whole-brain, network, and nodal changes compared to modularity and participation coefficient

To investigate whether entropy uniquely co-varied with age compared to two commonly used graph theory metrics, we examined entropy’s relationship with age in a linear regression controlling for modularity and participation coefficient as covariates. Both modularity R(713) = - 0.51, *p* = 6.69e-48 and whole-brain participation coefficient R(713) = 0.28, *p* = 5.51e-14 significantly correlated with age (Figure 3a, 3b; Supplementary Table 1, though the effect sizes were lower in the Schaefer atlas calculations (Supplementary Figure 3; Supplementary Table 5).

**Figure 3.**
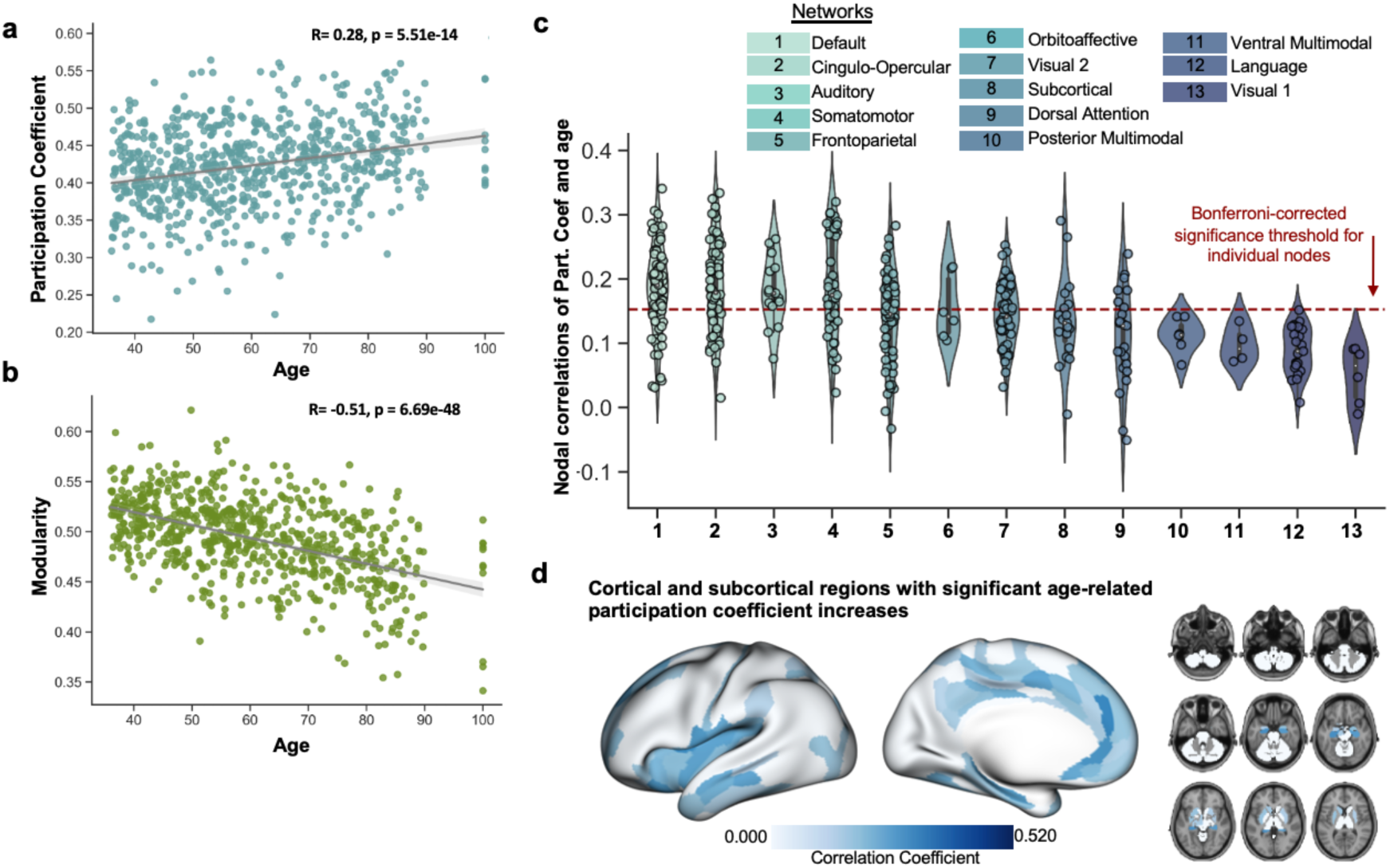
Whole-brain Modularity and Participation Coefficient Correlated with Age. **a.** Scatter plot of correlation between averaged whole-brain participation coefficient and age **b.** Scatter plot of correlation between averaged whole-brain modularity and age. **c.** Violin plot of nodal correlations between age and participation coefficient, organized by network. Red dotted line indicates Bonferroni-corrected significance threshold for nodal relationships, **d.** Brain plots of nodes with significant age-participation coefficient correlations.

When controlling for covariates, entropy’s relationship with age was the largest of the three with an effect size of 0.48 compared to 0.42 for modularity and 0.18 for participation coefficient (Table 2a, Supplementary Table 1). To determine the unique variance of entropy’s relationship with age, we included modularity and whole-brain averaged participation coefficient as additional covariates in the regression of entropy on age, finding that entropy’s relationship with age remained significant (*B* = 0.29, *p* = 8.36e-11) (Table 2a). Examining the relationship between participation coefficient and age at the nodal and network level, we found that, in contrast to entropy, only four networks demonstrated a majority of nodes with significant age-changes in the Glasser atlas (Figure 3c), and three networks in the Schaefer atlas (Supplementary Figure 3c). Participation Coefficient nodal age-correlations were also less widespread and weaker than entropy (Figure 2d, Supplementary Figure 3d).

### Entropy moderates cross-sectional age-related declines in fluid cognition

Next, we examined whether task-evoked whole-brain entropy moderates the relationship between increasing age and decreasing fluid cognition scores. As expected, we found that age was negatively correlated with fluid cognition scores as measured by the NIH Toolbox R(614)= - 0.5698, *p*<.001. Entropy significantly moderated this relationship, (*B* = −0.11, *t*(603) = −2.79, *p* = 5.04e-03; Table 2b), indicating that individuals with lower entropy demonstrated higher fluid cognition scores in old age. The relationship between age and cognition was significant for all values of entropy in this dataset, however, a simple slope analysis revealed that when entropy was one standard deviation below the mean, the relationship between age and fluid cognition was significantly less steep B = −0.39, *t*(603) = −8.36, *p* < 0.001 compared to when entropy was one standard deviation above the mean B = −0.56, *t*(603) = −13.52, *p* < 0.001(Figure 4). This relationship remained after controlling for covariates (Table 2b) and was similar in the Schaefer atlas (Supplementary Table 4). A similar moderating effect was found for modularity and participation coefficient (Supplementary Table 6), though the moderation effect of participation coefficient did not remain significant after controlling for demographic factors. Furthermore, Johnson-Neyman analysis demonstrated modularity’s overall significant moderation effect was in part driven by a significant moderating effect in young adults, whereas entropy’s moderation effect was specific to older adults.

**Figure 4.**
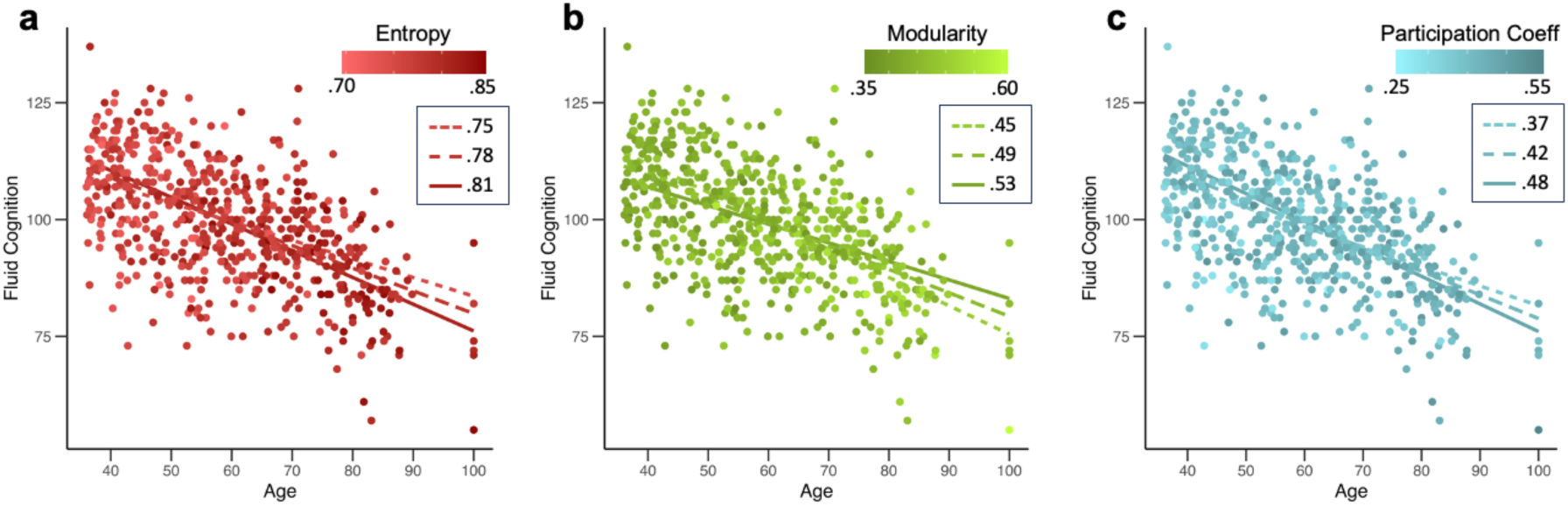
Entropy, Modularity, and Participation Coefficient as a Moderator of Age and Fluid Cognition Relationship. Relationship between age and fluid cognition for different values of **a.** entropy, **b.** modularity, and **c.** participation coefficient. Three lines represent the mean value and one standard deviation above and below the mean.

**Table 2b.** Linear interaction statistics between whole-brain task-averaged entropy and age on fluid cognition, accounting for covariates.

| Covariates | *b* | *SE* | *t* | *p* |
| --- | --- | --- | --- | --- |
| **Entropy\*Age** |  |  |  |  |
| No covariates | -0.11 | 0.04 | -2.79 | 5.04e-03** |
| **Accounting for in-scanner motion and brain volume** |  |  |  |  |
| MeanFD, MeanFD<sup>2</sup>, eTIV | -0.11 | 0.04 | -2.82 | 5.04e-03** |
| **Accounting for participant demographics, motion, brain volume** |  |  |  |  |
| Sex, Race, Ethnicity, Education, MeanFD, MeanFD<sup>2</sup>, eTIV | -0.08 | 0.03 | -2.38 | 0.02* |
| **Accounting for additional graph theory metrics, and all other covariates** |  |  |  |  |
| Participation Coef., Modularity, Sex, Race, Ethnicity, Education, MeanFD, MeanFD<sup>2</sup>, eTIV | -0.09 | 0.04 | -2.57 | 0.01* |
| **Accounting for graph theory metrics as moderators, and all other covariates** |  |  |  |  |
| Participation Coef.*age, Modularity*age, Participation Coef., Modularity, Sex, Race, Ethnicity, Education, MeanFD, MeanFD<sup>2</sup>, eTIV | -0.07 | 0.05 | -1.33 | 0.19 |

Notes. All graph theory metrics were calculated using functional connectivity matrices generated from surface-based cortical data applied to the Glasser atlas with volumetric subcortical data applied to the Cole atlas. Mean FD refers to the motion metric average framewise displacement. eTIV = estimated total intracranial volume.

### Entropy exhibits greater between-task reliability in age-related differences than modularity and participation coefficient

As an exploratory analysis, we examined task-specific relationships between age and entropy, modularity, and participation coefficient. We found that metrics measured during the attentional control task were the most strongly associated with age for modularity and participation coefficient, whereas the strongest relationship for entropy was from the episodic memory task (Supplementary Figure 7). Overall, whole-brain entropy demonstrated a medium effect size in its relationship with age for the attention, episodic memory, and processing speed tasks (*B* = 0.4, 0.45, 0.43), whereas modularity and participation coefficient showed more variability (Supplementary Table 3). For modularity, associations with age were large for the attention task and medium for processing speed and episodic memory (*B* = 0.52, 0.33, 0.29). For participation coefficient, the effect size of the relationship with age was medium for attention, small for episodic memory, and nonsignificant for processing speed (*B* = 0.35, 0.13, 0.06). To examine task-specific relationships with age for entropy, modularity and participation coefficient simultaneously, each metric’s task-specific score were entered into a regression on age. Entropy during the processing speed task (*B* = 0.13, *p* = 8.30e-04), entropy during the episodic memory task (*B* = 0.16, *p* = 2.04e-04), and modularity during the attentional task (*B* = −0.31, *p* = 8.69e-07) were significant unique correlates of age.

## Discussion

Previous examinations of functional network properties across the lifespan have provided converging findings on neural and network desegregation with advancing age^14–27^, which have been taken as functional connectivity correlates of neural dedifferentiation^25^. Despite this, there remain questions about whether we can use new methodologies and capitalize on task-based fMRI data to better understand processes of aging and age-related cognitive decline. The edge-centric framework employed in the current study allows us to address a few previous limitations. First, traditional methods for detecting modules generally force “hard partitions” of brain modules that are not biologically plausible and disallow nodes from participating in multiple modules simultaneously^31,33^. Second, only a few metrics are conceptually related to the neural dedifferentiation phenomenon, amenable to being investigated at the level of the whole brain, network, and individual node, and have been studied in aging^41^. The edge-centric approach, which results in overlapping communities, addresses these limitations by better conforming to the hypothesis that brain regions and systems can be activated for multiple unrelated functions ^33^. Our study focused on examining entropy, a metric representing the diversity with which nodes participated in these edge-defined communities, at the nodal, network, and whole-brain levels during three cognitive tasks (episodic memory, visual motor, and attention). Importantly, we found that entropy increases significantly across the whole brain with age while differentially increasing in specific regions and networks, giving rise to important questions on the origins of such entropic increases. Of note, we found that entropy did not significantly decrease in any nodes. Specifically, entropy significantly increased in 304 of the 379 nodes, suggesting a lack of compensatory decreased entropy in unaffected brain regions. Age-related changes to entropy were found to be regional but strong enough to still be detected by network and whole-brain measurements. Finally, we observed a wide range of entropy changes across the brain and demonstrated that lower entropy was associated with steeper increases in cognitive decline in older adults, suggesting entropy may be either a marker of pathological change or a marker of cognitive resilience.

A core observation in the study of brain aging has been the decline in recruitment and connectivity of specialized canonical networks with increasing age^42^. Older adults tend to recruit bilateral areas of the prefrontal cortex during tasks of attentional control^43^, episodic memory^44–46^, and executive functioning^45^, and the additional recruitment has been associated with both improved and deteriorated performance^45,46^. Functional connectivity studies further support this, demonstrating reduced intra-network connectivity, increased inter-network connectivity, and reduced modularity with advancing age^17,20^. Our study uses a large sample size from middle age to advanced age and provides support for the neural dedifferentiation theory of the aging brain. Across tasks spanning three cognitive domains impacted by aging—processing speed, episodic memory, and attentional control—we found evidence for increased entropy with advancing age, indicating that many nodes, networks, and the brain overall becomes less specialized. Given that we observed nodal entropy values ranging from 0.1 to 0.99 even in the youngest age range (ages 36-45), we speculate that high entropy values are not inherently atypical. Rather, we speculate that high entropy nodes are those whose participation in modules is more evenly divided among modules, which in younger adults may denote regions involved in the coordination of networks or facilitation of communication, and in older adults may denote either compensation or disorder.

A distinct advantage of the edge-centric approach is the ability to calculate a metric of specificity at the regional level, allowing us to examine how entropy increases with age at multiple levels of the brain’s hierarchical organization. We found that age-related changes in entropy primarily affected individual nodes and were spatially clustered rather than clustered by network. Notably, the strongest age-related changes were observed in regions shown to be implicated in classic activity-based neural dedifferentiation studies—medial temporal cortices, insular cortices, visual systems, and several subcortical regions ^47^. Additionally, we found the most consistent age-related entropy increases across tasks in the medial temporal lobe, including parts of the hippocampus, parahippocampal regions, and auditory cortices, regions that are also known to demonstrate age-related structural and functional changes and are critically implicated in Alzheimer’s disease^48–51^. Although speculative in the absence of positron emission tomography (PET) data, it is possible that the task-independent increased entropy in these transmodal regions, signifying the increased metabolic activity of these nodes, may potentially be one reason for the accumulation of neurodegenerative pathologies, such as beta-amyloid and tau. Previous work has shown that these pathologies spread along functional networks^52,53^ and participation of these individual nodes in multiple communities, particularly those involved in emergent cognitive functioning, could potentially lead to the aggregation of toxic proteinopathies. Conversely, it is also possible that the aggregation of amyloid and tau in these regions reduces specialized processing, increasing the entropy of these individual nodes. Future longitudinal, PET-based studies could help better understand the correspondence between changes in brain dynamics and the pathophysiological deposition of amyloid and tau.

Alternatively, these regions also exhibit significant cortical thinning throughout Alzheimer’s disease ^54^. Previous research has shown that age-related cortical thinning is most notable and consistent in frontal and parietal regions and less so in limbic and sensory regions ^55^, which we did not find to be consistently related to entropy in our study. We also controlled for total brain volume in these analyses, suggesting that entropy may not simply be a functional correlate of neurodegeneration. Future research examining mechanisms of age-related declines in nodal entropy, and how nodal changes affect network and whole-brain organization, may further inform our understanding of how pathological and normative aging processes result in cognitive and behavioral change.

It is important to note that no individual nodes exhibited a significant decrease in entropy with age, which supports previous research that nodes become less specialized with age^25,27^, and even brain regions that may be spared from significant pathology in healthy aging do not demonstrate a compensatory decrease in entropy. As previously mentioned, high entropy nodes were found in the youngest age group, and it is conceivable that some nodes could have displayed a compensatory pattern of decreasing entropy in aging. In brain development, environmental factors such as socio-economic status and the richness of experiences have been linked to network segregation trajectories^56^. Thus, one possible explanation for these findings is that network segregation must be optimized during development and may be prevented from declining. However, age-related compensatory reorganization does not entail ‘healthier’ brain regions increasing nodal specialization to compensate for de-specialization elsewhere.

We also provide evidence for entropy as a unique neural correlate of age. Entropy is a metric used in information theory to investigate the pattern of communication within a system, where high entropy can indicate a system with greater or randomness in its communication patterns^35^. In a biological system such as the human brain, we can apply entropy to a distribution of nodal participation in overlapping communities to establish the degree to which a node’s connectivity with other nodes is specialized to a rare few or spread randomly among potential connectivity partners^33^. Given that biological systems often organize in specialized sub-units^57^, reduced entropy may indicate the brain’s failure to exert or maintain this pressure, which is not captured by existing metrics. As such, we compared entropy to two traditional graph theory metrics: whole-brain modularity and nodal participation coefficient, which through prior research are known to decline with advancing age^15,1758^. We found that whole brain entropy and modularity exhibited statistical overlap but were uniquely correlated with age. Furthermore, at the whole-brain, network, and nodal level, entropy demonstrated stronger and more consistent age-related changes than modularity and participation coefficient. Nodal participation demonstrated weaker and less consistent age-related and cognitive changes compared to entropy, which is consistent with previous findings that some node-based graph theory measures show variable age effects dependent on the specific in-scanner tasks employed in the study^14^.

A secondary objective of this study was to examine the role of entropy in the relationship between aging and cognition. We found that whole-brain entropy moderated the relationship between age and fluid cognition. Theoretical models of cognitive aging have previously acknowledged the possibility of this moderating relationship^59^, however much of the work in this field has favored examination of the possible mechanistic role^60–62^ It is important to note that these two pathways—moderation and mediation—are not mutually exclusive. Rather, metrics such as entropy and modularity are measured in a complex system and are likely both affected by age-related processes (mediation) and, at measurement time, reflect the degree of network resilience and therefore impact of age-related processes on cognition (moderation). However, in this study, we did not observe a significant direct relationship between global network properties and cognition. One potential interpretation of entropy’s increase with age is that nodes de-specialize as either a mechanism for or as a result of compensation for maintained complex behavior. In the current study, our observation of lower whole-brain entropy was indicative of better cognitive function across the lifespan suggesting that entropy increases are potentially a marker of nodal health rather than compensation. Furthermore, our preliminary moderation probing suggests that the relationship between network segregation and cognition may differ across the lifespan, with nodal specialization and network segregation becoming more important for cognition in old age. Previous work has characterized whole-brain modularity as a biomarker of neuroplasticity ^37^, thus, future work is needed to investigate whether metrics such as entropy and modularity reflect a protective factor that decreases with age or are better conceptualized as a neuromarker of damage accumulation.

The results of this study are limited by the fact that this data was collected cross-sectionally, and data collection was limited to healthy adults. Thus, this sample is subject to both cohort effects and a sampling bias that becomes more prevalent in older adults. Additionally, individuals in this study, particularly those in the oldest-old age group, are less representative of ‘typical aging’ than studies that include individuals with chronic health conditions or symptoms of cognitive impairment. Thus, future work should examine the relationships between entropy, age, and cognition in participant groups that include patient populations. This may also help bridge understanding of mechanisms of entropy increases with age, given that these increases may be more significant or widespread in individuals with neurological disease. Another limitation is our use of relatively short durations of task-based fMRI data. Previous work has examined entropy in resting state fMRI and during movie watching for longer periods of time^33^. For this reason, we examined entropy values averaged across tasks and examined consistency between tasks as a supplementary analysis. Between tasks, we found that entropy appeared to be more consistently associated with age than modularity or participation coefficient. It has also recently been proposed that use of task data in functional connectivity studies is subject to a potentially confounding role of activation-based signal in functional connectivity estimates^63^. While we did not explore this in the current study given our primary objective of establishing entropy within a literature that has not historically controlled for this, we offer that future studies might attempt to disentangle these effects in both edge-based and node-based connectivity approaches.

In conclusion, the current study found that entropy is a robust and unique neural correlate of aging with potentially important implications for cognitive health. We found that entropy increases present differentially across the whole brain, strongest and most consistent in medial-temporal, insular and subcortical regions, with strong enough age effects to still be detected when averaged at the canonical network and at the whole-brain levels. Importantly, we found evidence that the negative relationship between age and fluid cognition is moderated by entropy and modularity. Together, these findings support entropy as a potentially important metric to examine how pathological processes associated with aging, as well as interventions and lifestyle choices, affect functional neural dedifferentiation at the nodal, network, and whole-brain level. Furthermore, edge-connectivity entropy is a conceptually unique metric for the study of age-related changes. When used in conjunction with network segregation measures, we may be able to differentiate the importance of these two facets of network change in explaining variance in behavioral and pathological processes. The current study suggests that aging significantly and uniquely influences edge-based and node-based connectivity measures but that cognitive changes may be captured by either of these approaches similarly.

## Methods

### Participants

This study utilized data from the Lifespan Human Connectome Project Aging (HCP-A) 2.0 Release, containing 725 unrelated healthy individuals between the ages of 34 and 100+ years old (55.7% female). This study aimed to focus on the ‘typical’ aging process^64^, participants whose health is typical for their age. As such, participants include individuals with prevalent health conditions, including hypertension, but excludes individuals with less prevalent conditions such as diagnosed or suspected Alzheimer’s Disease and symptomatic stroke, as well as individuals with macular degeneration or poor cognitive performance for their age as measured by the MoCA and the TICS-M^64^.

#### NIH Cognitive Toolbox

Fluid cognition scores were calculated from five sub-domains of cognitive performance tested by the NIH Cognitive Toolbox, which has been shown to be reliable, valid, and concordant with standardized neuropsychological measures of cognition in healthy and clinical populations^65–67^. These cognitive measures consisted of the Dimensional Change Card Sort Test, the Flanker Inhibitory Control and Attention Test, the Picture Sequence Memory Test, and the Pattern Comparison Processing Speed Test. Due to our interest in entropy as a moderating factor of the established negative relationship between age and cognition, we utilized age-uncorrected standard scores.

### Neuroimaging

#### Imaging Protocols

A detailed account of the imaging protocols and MRI acquisition can be found in the imaging protocol for the HCP-A dataset^68^. The HCP-A is a multi-site study; all scanning sites used a Siemens 3T Prisma scanner and a 32-channel head enabling multiband data acquisition. Participants completed two, 45-minute imaging sessions as a part of their participation in the study, during which T1w and T2w structural images were acquired. As part of the functional MRI collection, in their first session, participants complete three fMRI scans collected during the performance of a cognitive task: a “Go/NoGo” attentional task, a “Vismotor” visuomotor processing speed task, and a “FaceName” episodic memory task. In the Go/NoGo task participants are instructed to respond with a button press as quickly as possible when any shape appears on the screen unless it is one of two target shapes, in which case they are asked to inhibit their response. In the Vismotor task, participants are instructed to observe a black and white circular checkerboard and respond as quickly as possible with a left button press with their index finger or a right button press with their middle finger when they see a red flickering square in the left or right half of the circle. In the FaceName task, participants are instructed to memorize pairs of names and face, retain these associations during Go/NoGo distractor trials, and then recall names in response to being cued with the previously seen faces.

#### Preprocessing

This study utilized a subset of the HCP-A data release pre-selected and minimally preprocessed by the HCP study coordinators (see processing pipeline protocol^69^ for detailed preprocessing procedure). Briefly, PreFreeSurfer was used to align T1w and T2w images, perform bias field correction and register the participant’s native structural volume space to MNI space. FreeSurfer was used to segment the volumes into predefined structures, reconstruct the white and pial cortical surfaces, and perform surface-based registration. PostFreeSurfer was used to produce a final brain mask, cortical surface files, and volumetric sub-cortical files. For the current study we utilized CIFTI files containing cortical data in grayordinate space and subcortical data in volumetric space. These data were minimally preprocessed by the HCP coordinators, including surface smoothing with a 2mm gaussian kernel and application of an HCP-specific ICA-Fix pipeline for noise removal. This pipeline is well validated for noise removal in the HCP datasets^70^, however it does not remove global signal, which has been demonstrated to additionally improve physiological noise reduction in HCP data^71^. Here, we averaged the signal over all cortical grayordinates and regressed this signal from the data using MATLAB. The resulting data was parcellated using the Glasser Atlas^72^, resulting in 360 cortical regions and 19 subcortical regions.

### Network Construction and Graph Theory Measures

#### Functional Connectivity Matrices, Modularity Maximization, and Participation Coefficient

Functional connectivity matrices were calculated by Pearson correlating the timeseries signal from each node in the Glasser atlas with every other node, resulting in a 379×379 symmetrical matrix with each cell representing the average correlation of two nodes. Whole-brain modularity was calculated using the Community-Louvain technique using the Brain Connectivity Toolbox (BCT)^10^ using weighted, unthresholded connectivity matrices and applying an asymmetric treatment of negative weights^10,73^. This algorithm segregates nodes into distinct, non-overlapping communities (modules) by iteratively assigning nodes to modules that maximize the value of the overall modularity factor, Q^74^. Given that this algorithm may vary depending on the random order of node selection, this process was repeated 1000 times per individual to create a distribution of module assignments for each node for each individual. From this, an agreement matrix was calculated containing the proportion of time two nodes are in the same module. This was used as an input into a consensus clustering technique^75^ to create a consensus community partition, which uses the Community-Louvain algorithm to maximize Q. This was repeated 1000 times to create a new agreement matrix until the modules converge to a single clustering. To calculate a stable modularity value of Q, we input this converged module assignment matrix into the Community-Louvain algorithm with the participant’s weighted connectivity matrix to calculate their final maximal modularity value^76^. The converged module assignment matrix was then used to calculate participation coefficient for each node with the BCT function “participation_coef_sign.m”.

#### Edge Timeseries Construction and Entropy Calculation

Construction of the edge time series consisted of taking the element-wise product of the z-scored parcel timeseries for every pair of nodes in the connectivity matrix^33^. This results in a N x T size matrix where N equals the number of nodes in the parcellation scheme and T equals the number of temporal observations. Each instance of T in this matrix reflects the co-fluctuation amplitude of two nodes at a given time point in the functional run, allowing for the retention of temporal information in the relationship between nodes. Entropy was calculated by applying a k-means clustering algorithm to the edge timeseries data, which assigns a community label to each edge. From this, the distribution of each node’s participation in the identified edge communities is calculated. Entropy is then calculated as:

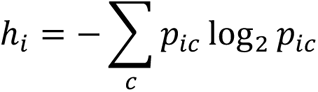

Where *h_i_* is entropy of the *i*th node and *p_ic_* is fraction of the *i*th node’s connections assigned to edge community *c*. Entropy calculated at the level of the individual node allows for examination of entropy at multiple levels of the functional hierarchy, and so to examine and capitalize on this we also calculated average entropy scores for regions of the brain involved in canonical networks and the whole brain.

### Statistics

All statistics were performed in MATLAB version 2022a, with the exception of the moderation probing analyses for the second aim, which were performed in R 2022.02.2 using the “interactions” toolbox.

