## Supplementary Materials for "Edge Community Entropy is a Novel Neural Correlate of Aging and Moderator of Fluid Cognition"

**Supplementary Table 1| Linear regression statistics of relationship between whole-brain task-averaged modularity and participation coefficient and age, accounting for covariates**

| Covariates | *b* | *SE* | *t* | *p* |
|---|---|---|---|---|
| **Modularity** | | | | |
| No covariates | -0.50 | 0.03 | -15.70 | 6.69e-48*** |
| **Accounting for motion, brain volume** | | | | |
| MeanFD, MeanFD$^2$, eTIV | -0.43 | 0.03 | -12.71 | 1.74e-33*** |
| **Accounting for participant demographics, motion, brain volume** | | | | |
| Sex, Race, Ethnicity, Education, MeanFD, MeanFD$^2$, eTIV | -0.42 | 0.03 | -12.75 | 1.3e-33*** |
| **Accounting for additional graph theory metrics, and all other covariates** | | | | |
| Participation Coef., Entropy, Sex, Race, Ethnicity, Education, MeanFD, MeanFD$^2$, eTIV | -0.38 | 0.06 | -6.12 | 1.54e-09*** |
| **Participation Coef.** | | | | |
| No covariates | 0.27 | 0.04 | 7.67 | 5.51e-14*** |
| **Accounting for motion, brain volume** | | | | |
| MeanFD, MeanFD$^2$, eTIV | 0.20 | 0.03 | 6.00 | 3.08e-09*** |
| **Accounting for participant demographics, motion, brain volume** | | | | |
| Sex, Race, Ethnicity, Education, MeanFD, MeanFD$^2$, eTIV | 0.18 | 0.03 | 5.29 | 1.61e-07*** |
| **Accounting for additional graph theory metrics, and all other covariates** | | | | |
| Entropy, Modularity, Sex, Race, Ethnicity, Education, MeanFD, MeanFD$^2$, eTIV | -0.19 | 0.04 | -4.15 | 3.82e-05*** |

Notes. All graph theory metrics were calculated using functional connectivity matrices generated from surface-based cortical data applied to the Glasser atlas with volumetric subcortical data applied to the Cole atlas. Mean FD refers to the motion metric average framewise displacement. eTIV = estimated total intracranial volume.

**Supplementary Table 2| Linear interaction statistics between whole-brain task-averaged modularity and participation coefficient and age on fluid cognition, accounting for covariates**

| Covariates | *b* | *SE* | *t* | *p* |
|---|---|---|---|---|
| **Modularity** | | | | |
| No covariates | 0.13 | 0.04 | 3.49 | 5.27e-04*** |
| **Accounting for in-scanner motion and brain volume** | | | | |
| MeanFD, MeanFD$^2$, eTIV | 0.12 | 0.04 | 3.34 | 8.83e-04*** |
| **Accounting for participant demographics, motion, brain volume** | | | | |
| Sex, Race, Ethnicity, Education, MeanFD, MeanFD$^2$, eTIV | 0.08 | 0.03 | 2.38 | 0.02* |
| **Accounting for additional graph theory metrics, and all other covariates** | | | | |
| Participation Coef., Modularity, Sex, Race, Ethnicity, Education, MeanFD, MeanFD$^2$, eTIV | 0.08 | 0.03 | 2.34 | 0.02* |
| **Accounting for graph theory metrics as moderators, and all other covariates** | | | | |
| Participation Coef.*age, Modularity*age, Participation Coef., Modularity, Sex, Race, Ethnicity, Education, MeanFD, MeanFD$^2$, eTIV | 0.02 | 0.07 | 0.35 | 0.73 |
| **Participation Coef.** | | | | |
| No covariates | -0.11 | 0.04 | -2.90 | 3.90e-03** |
| **Accounting for in-scanner motion and brain volume** | | | | |
| MeanFD, MeanFD$^2$, eTIV | -0.10 | 0.04 | -2.75 | 6.12e-03** |
| **Accounting for participant demographics, motion, brain volume** | | | | |
| Sex, Race, Ethnicity, Education, MeanFD, MeanFD$^2$, eTIV | -0.05 | 0.03 | -1.63 | 0.10 |
| **Accounting for additional graph theory metrics, and all other covariates** | | | | |
| Participation Coef., Modularity, Sex, Race, Ethnicity, Education, MeanFD, MeanFD$^2$, eTIV | -0.06 | 0.03 | -1.70 | 0.09 |
| **Accounting for graph theory metrics as moderators, and all other covariates** | | | | |
| Participation Coef.*age, Modularity*age, Participation Coef., Modularity, Sex, Race, Ethnicity, Education, MeanFD, MeanFD$^2$, eTIV | -0.01 | 0.05 | -0.27 | 0.79 |

Notes. All graph theory metrics were calculated using functional connectivity matrices generated from surface-based cortical data applied to the Glasser atlas with volumetric subcortical data applied to the Cole atlas. Mean FD refers to the motion metric average framewise displacement. eTIV = estimated total intracranial volume.

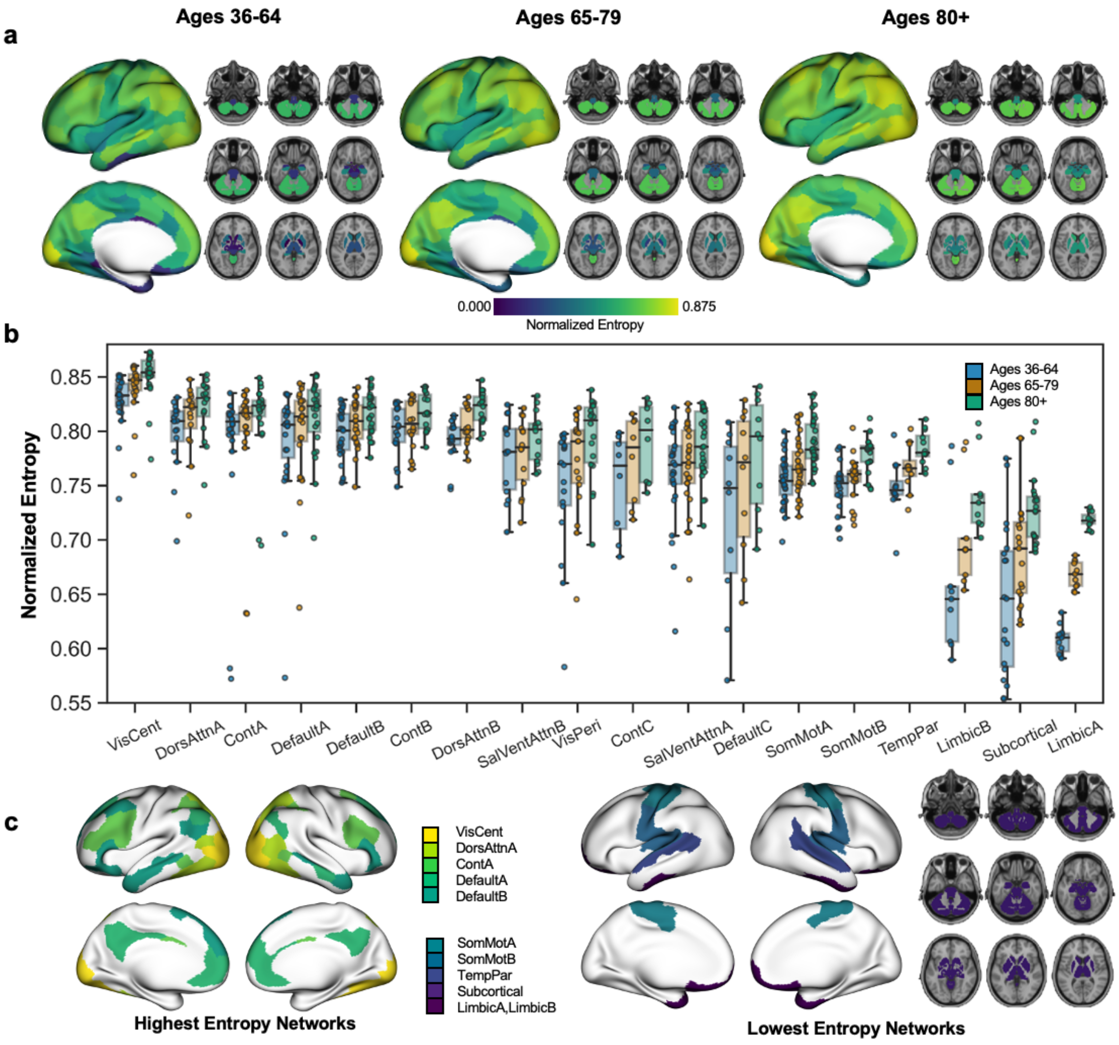

**Supplementary Figure 1. Normalized Entropy using Schaefer Atlas. a.** Normalized entropy at the nodal level across the whole brain, averaged across HCPA-defined age groups of 36-64, 65-79, and 80-100+. **b.** Normalized entropy at the network level by HCPA-defined age groups. **c.** Networks with the highest normalized entropy values across the entire age range (The Visual Central, Dorsal Attention A, Control A, Default A, and Default B networks). **d.** Networks with the lowest normalized entropy values across the entire age range (The Somatomotor A & B, Tempoparietal, Limbic A & B networks, and subcortical regions).

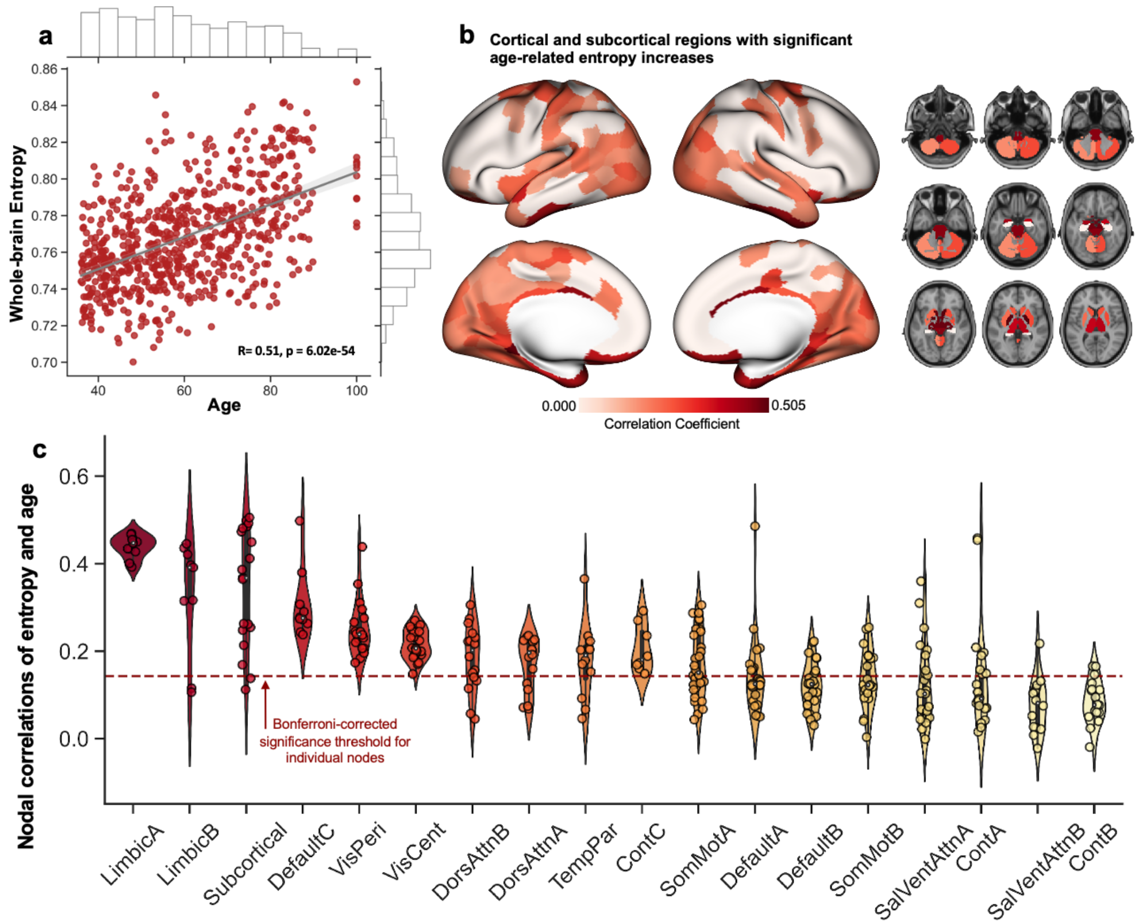

**Supplementary Figure 2. Entropy correlated with age using Schaefer Atlas. a.** Scatter plot of correlation between averaged whole-brain entropy and age. **b.** Brain plots of nodes with significant age-entropy correlations. **c.** Violin plot of nodal correlations between age and entropy organized by network. Red dotted line indicates Bonferroni-corrected significance threshold for nodal relationships.

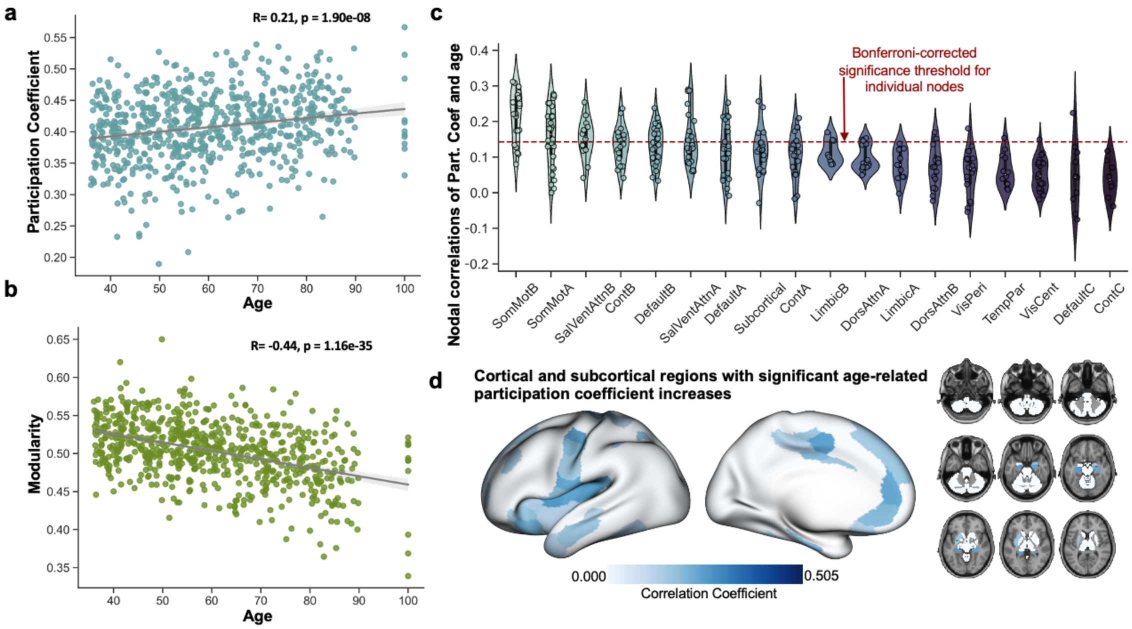

**Supplementary Figure 3. Whole-brain Modularity and Participation Coefficient using Schaefer Atlas.**
**a.** Scatter plot of correlation between averaged whole-brain participation coefficient and age **b.** Scatter plot of correlation between averaged whole-brain modularity and age. **c.** Violin plot of nodal correlations between age and participation coefficient, organized by network. Red dotted line indicates Bonferroni-corrected significance threshold for nodal relationships. **d.** Brain plots of nodes with significant age-participation coefficient correlations.

**Supplementary Table 3| Individual Linear Model Results for Attentional, Episodic Memory, and Processing Speed Task Graph Theory Metrics**

| | *b* | *p* |
|---|---|---|
| **Entropy, No covariates** | | |
| Attention | 0.40 | 6.23e-30 |
| Episodic Memory | 0.45 | 4.14e-37 |
| Processing Speed | 0.43 | 2.56e-33 |
| **Modularity, No covariates** | | |
| Attention | -0.52 | 1.29e-52 |
| Episodic Memory | -0.33 | 5.5e-20 |
| Processing Speed | -0.29 | 1.84e-15 |
| **Participation Coef., No covariates** | | |
| Attention | 0.35 | 1.50e-22 |
| Episodic Memory | 0.13 | 4.71e-04 |
| Processing Speed | 0.06 | 0.14 |

Notes. All graph theory metrics were calculated using functional connectivity matrices generated from surface-based cortical data applied to the Glasser atlas with volumetric subcortical data applied to the Cole atlas.

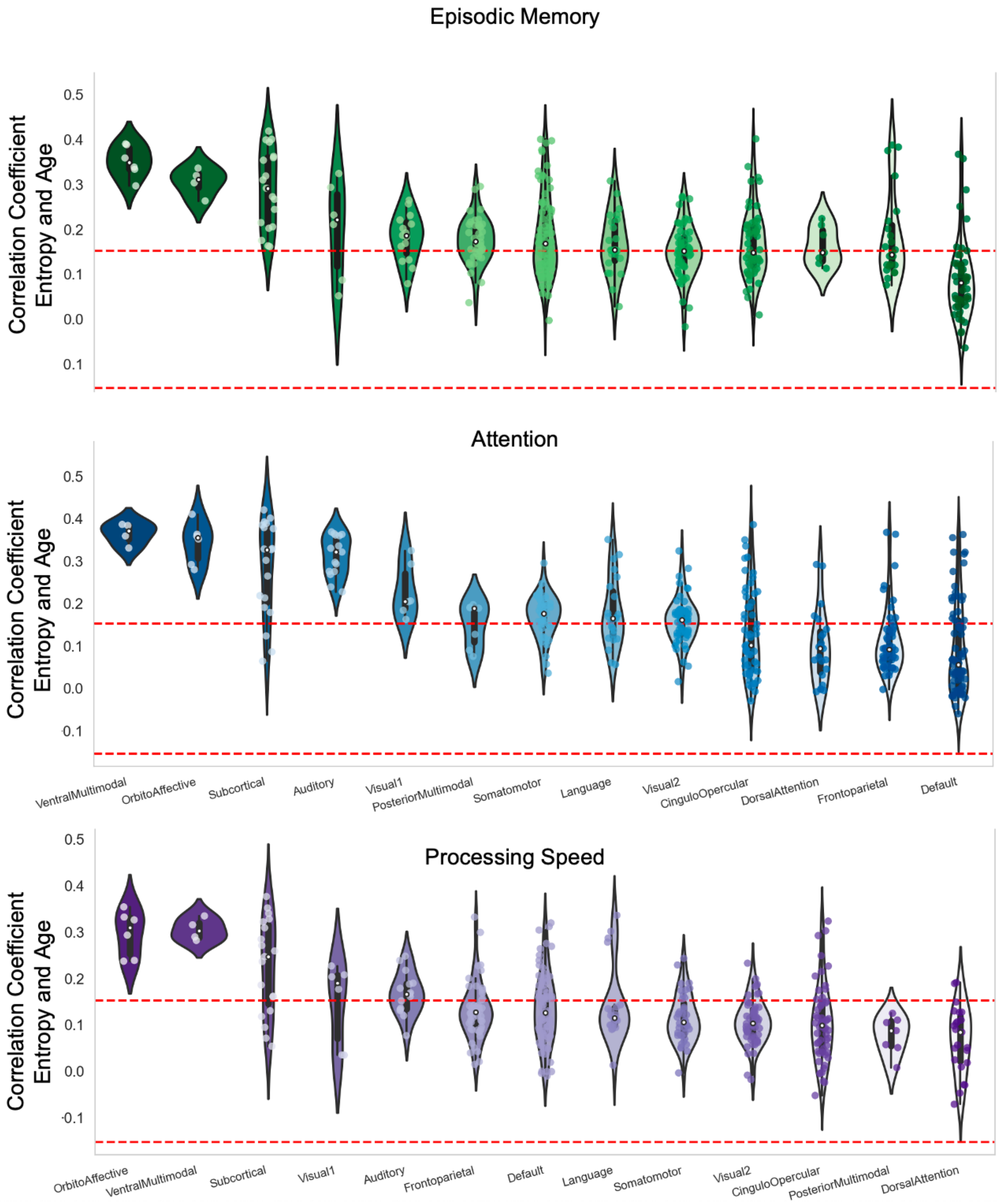

**Supplementary Figure 4. Correlation coefficients for the relationship between age and entropy for individual tasks, grouped by Glasser atlas networks.** Regions with significant age-related increases in entropy across the lifespan, for the attentional task, episodic memory task, and processing speed task. Red lines indicate Bonferroni significance thresholds for nodal-level correlations, corrected for 1137 comparisons (3 tasks, 379 nodes).

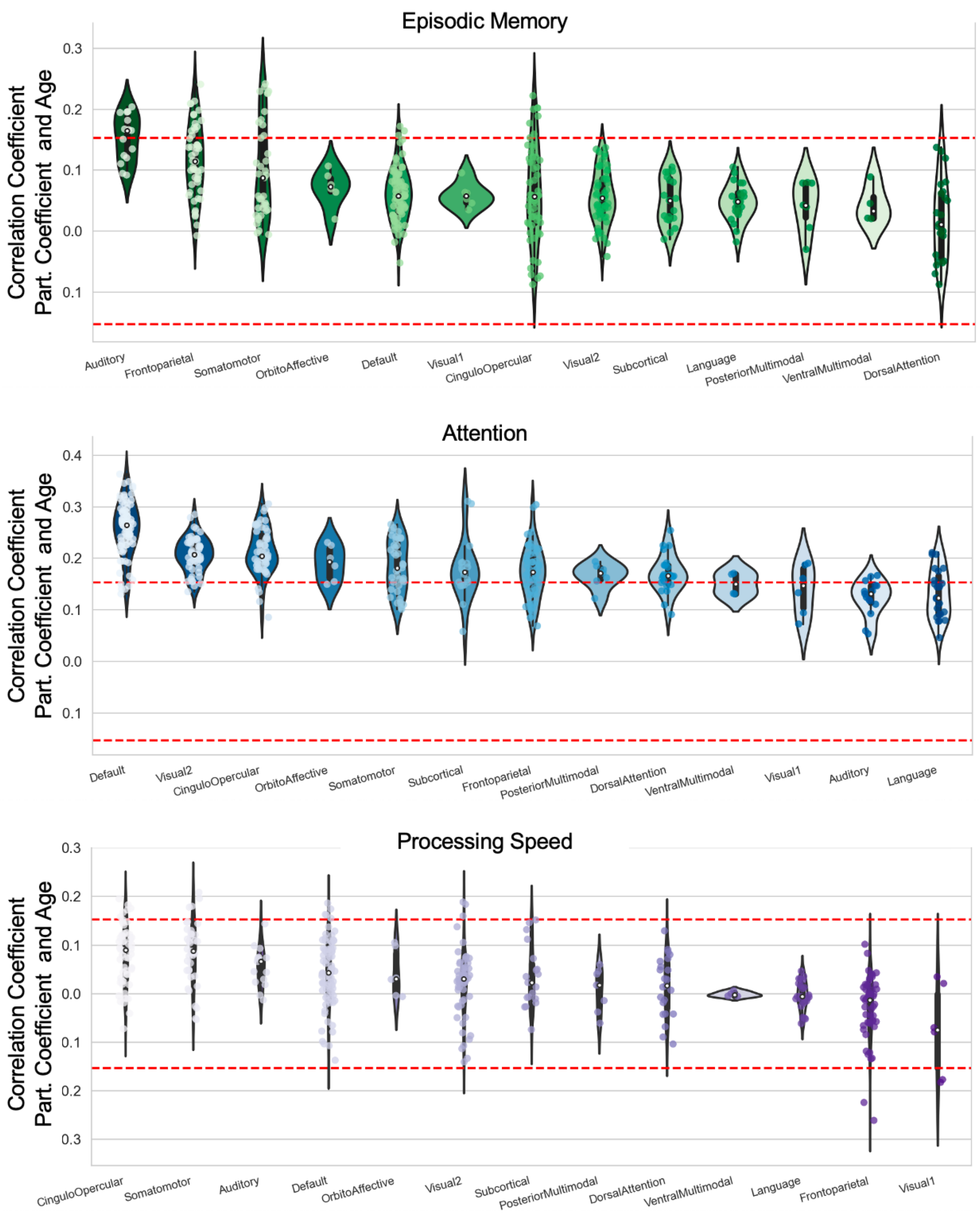

**Supplementary Figure 5. Correlation coefficients for the relationship between age and participation coefficient for individual tasks, grouped by Glasser atlas networks.** Regions with significant age-related increases in entropy across the lifespan for the attentional task, episodic memory task, and processing speed task. Red lines indicate Bonferroni significance thresholds for nodal-level correlations, corrected for 1137 comparisons (3 tasks, 379 nodes).

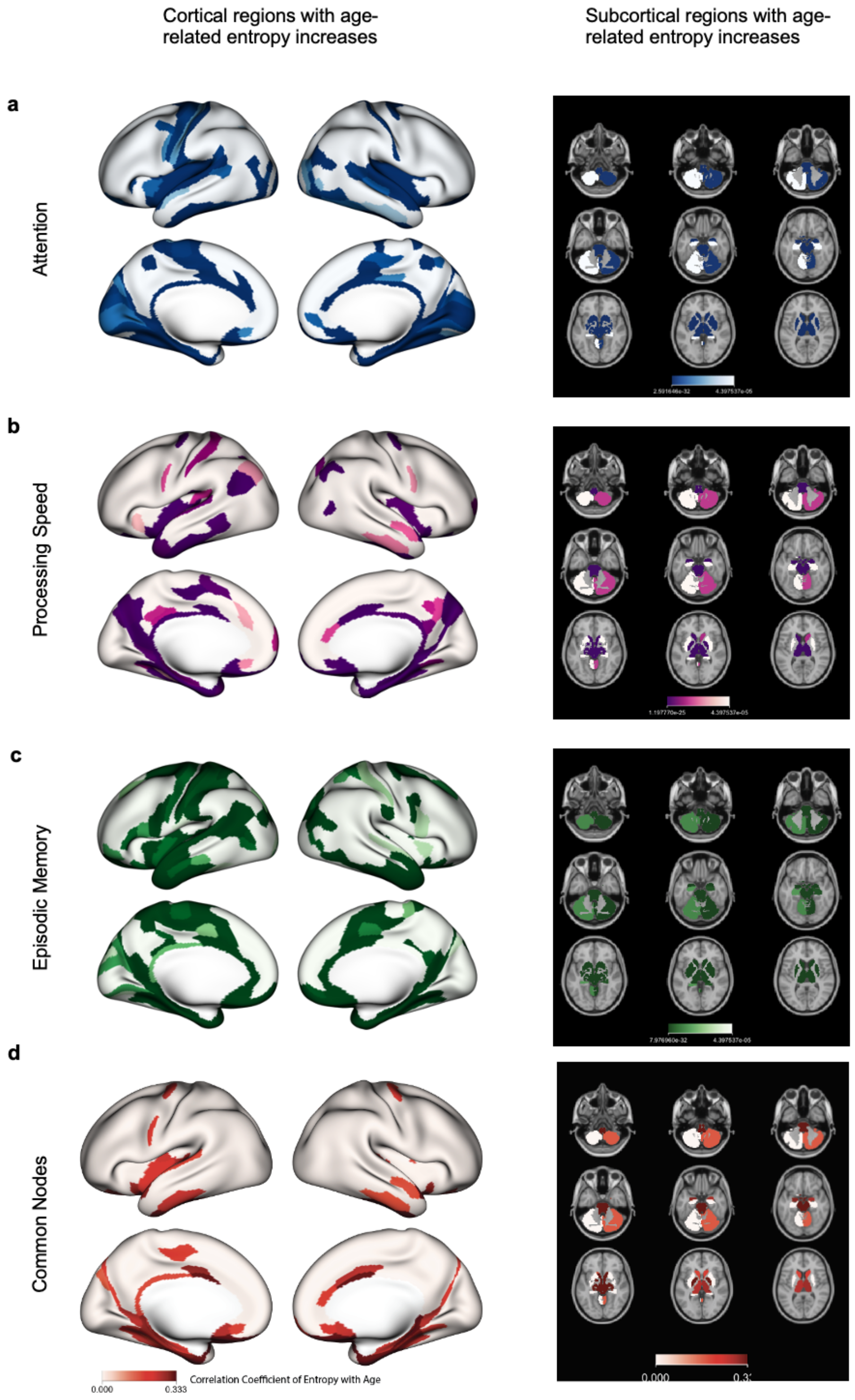

**Supplementary Figure 6. Nodes with significant age-related entropy increases across individual tasks.** Regions with significant age-related increases in entropy across the lifespan, Bonferroni-corrected, for the **a**. attentional task, **b.** processing speed task, **c.** episodic memory task, **d.** and a representation of the overlap across these three tasks, representing the most consistent age-related entropy increases.

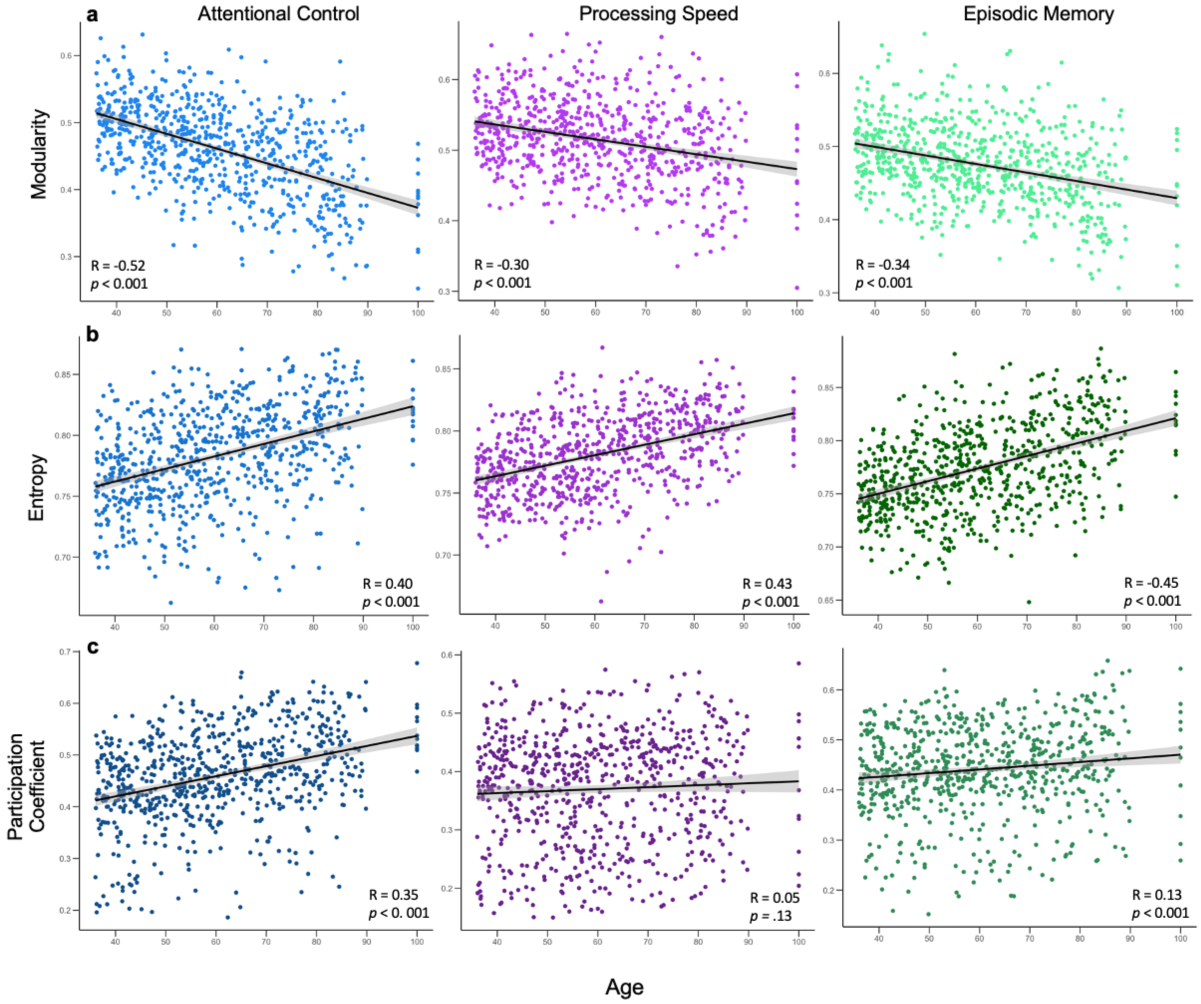

**Supplementary Figure 7. Whole-brain entropy, modularity and participation coefficient correlated with age for each cognitive task. a.** Scatter plot of correlation between averaged whole-brain modularity and age for the attentional control, processing speed, and episodic memory tasks. **b.** Scatter plot of correlation between averaged whole-brain entropy and age for the attentional control, processing speed, and episodic memory tasks. **c.** Scatter plot of correlation between averaged whole-brain participation coefficient and age for the attentional control, processing speed, and episodic memory tasks.

**Supplementary Table 4a| Linear regression statistics of relationship between whole-brain task-averaged entropy and age, accounting for covariates**

| Covariates | *b* | *SE* | *t* | *p* |
|---|---|---|---|---|
| **Entropy** | | | | |
| No covariates | 0.51 | 0.03 | 15.79 | 2.23e-48*** |
| **Accounting for motion, brain volume** | | | | |
| MeanFD, $MeanFD^2$, eTIV | 0.45 | 0.03 | 13.25 | 6.34e-36*** |
| **Accounting for participant demographics, motion, brain volume** | | | | |
| Sex, Race, Ethnicity, Education, MeanFD, $MeanFD^2$, eTIV | 0.45 | 0.03 | 13.71 | 5.51e-38*** |
| **Accounting for additional graph theory metrics, and all other covariates** | | | | |
| Participation Coef., Modularity, Sex, Race, Ethnicity, Education, MeanFD, $MeanFD^2$, eTIV | -0.26 | 0.05 | -5.17 | 3.01e-07*** |

**Supplementary Table 4b| Linear interaction statistics between whole-brain task-averaged entropy and age on fluid cognition, accounting for covariates**

| Covariates | *b* | *SE* | *t* | *p* |
|---|---|---|---|---|
| **Entropy*Age** | | | | |
| No covariates | -0.10 | 0.04 | -2.64 | 8.38e-03** |
| **Accounting for in-scanner motion and brain volume** | | | | |
| MeanFD, $MeanFD^2$, eTIV | -0.10 | 0.04 | -2.90 | 4.28e-03** |
| **Accounting for participant demographics, motion, brain volume** | | | | |
| Sex, Race, Ethnicity, Education, MeanFD, $MeanFD^2$, eTIV | -0.08 | 0.03 | -2.38 | 0.02* |
| **Accounting for additional graph theory metrics, and all other covariates** | | | | |
| Participation Coef., Modularity, Sex, Race, Ethnicity, Education, MeanFD, $MeanFD^2$, eTIV | -0.09 | 0.03 | -2.56 | 0.01* |
| **Accounting for graph theory metrics as moderators, and all other covariates** | | | | |
| Participation Coef.*age, Modularity*age, Participation Coef., Modularity, Sex, Race, Ethnicity, Education, MeanFD, $MeanFD^2$, eTIV | -0.08 | 0.05 | -1.80 | 0.07 |

Notes. All graph theory metrics were calculated using functional connectivity matrices generated from surface-based cortical data applied to the Schaefer atlas with volumetric subcortical data applied to the Cole atlas. Mean FD refers to the motion metric average framewise displacement. eTIV = estimated total intracranial volume.

**Supplementary Table 5 | Linear regression statistics of relationship between whole-brain task-averaged modularity and participation coefficient and age, accounting for covariates**

| Covariates | *b* | *SE* | *t* | *p* |
|---|---|---|---|---|
| **Modularity** | | | | |
| No covariates | -0.44 | 0.03 | -13.18 | 1.16e-35 |
| **Accounting for motion, brain volume** | | | | |
| MeanFD, MeanFD$^2$, eTIV | -0.36 | 0.03 | -10.85 | 1.86e-25 |
| **Accounting for participant demographics, motion, brain volume** | | | | |
| Sex, Race, Ethnicity, Education, MeanFD, MeanFD$^2$, eTIV | -0.35 | 0.03 | -10.59 | 2.20e-24 |
| **Accounting for additional graph theory metrics, and all other covariates** | | | | |
| Participation Coef., Entropy, Sex, Race, Ethnicity, Education, MeanFD, MeanFD$^2$, eTIV | -0.39 | 0.05 | -6.16 | 1.24e-07 |
| **Participation Coef.** | | | | |
| No covariates | 0.21 | 0.04 | 5.69 | 1.90e-08*** |
| **Accounting for motion, brain volume** | | | | |
| MeanFD, MeanFD$^2$, eTIV | 0.15 | 0.03 | 4.36 | 1.53e-05*** |
| **Accounting for participant demographics, motion, brain volume** | | | | |
| Sex, Race, Ethnicity, Education, MeanFD, MeanFD$^2$, eTIV | 0.12 | 0.03 | 3.58 | 3.72e-04*** |
| **Accounting for additional graph theory metrics, and all other covariates** | | | | |
| Entropy, Modularity, Sex, Race, Ethnicity, Education, MeanFD, MeanFD$^2$, eTIV | -0.26 | 0.05 | -5.17 | 3.01e-07*** |

Notes. All graph theory metrics were calculated using functional connectivity matrices generated from surface-based cortical data applied to the Schaefer atlas with volumetric subcortical data applied to the Cole atlas. Mean FD refers to the motion metric average framewise displacement. eTIV = estimated total intracranial volume.

**Supplementary Table 6 | Linear interaction statistics between whole-brain task-averaged modularity and participation coefficient and age on fluid cognition, accounting for covariates**

| Covariates | *b* | *SE* | *t* | *p* |
|---|---|---|---|---|
| **Modularity** | | | | |
| No covariates | 0.14 | 0.04 | 3.81 | 1.51e-04*** |
| **Accounting for in-scanner motion and brain volume** | | | | |
| MeanFD, $MeanFD^2$, eTIV | 0.12 | 0.04 | 3.42 | 6.77e-04*** |
| **Accounting for participant demographics, motion, brain volume** | | | | |
| Sex, Race, Ethnicity, Education, MeanFD, $MeanFD^2$, eTIV | 0.07 | 0.03 | 2.11 | 0.03* |
| **Accounting for additional graph theory metrics, and all other covariates** | | | | |
| Participation Coef., Modularity, Sex, Race, Ethnicity, Education, MeanFD, $MeanFD^2$, eTIV | 0.07 | 0.03 | 2.06 | 0.04* |
| **Accounting for graph theory metrics as moderators, and all other covariates** | | | | |
| Participation Coef.*age, Modularity*age, Participation Coef., Modularity, Sex, Race, Ethnicity, Education, MeanFD, $MeanFD^2$, eTIV | -0.02 | 0.07 | -0.36 | 0.72 |
| **Participation Coef.** | | | | |
| No covariates | -0.12 | 0.04 | -3.27 | 1.11e-03** |
| **Accounting for in-scanner motion and brain volume** | | | | |
| MeanFD, $MeanFD^2$, eTIV | -0.11 | 0.04 | -2.94 | 3.36e-03** |
| **Accounting for participant demographics, motion, brain volume** | | | | |
| Sex, Race, Ethnicity, Education, MeanFD, $MeanFD^2$, eTIV | -0.06 | 0.03 | -1.84 | 0.07 |
| **Accounting for additional graph theory metrics, and all other covariates** | | | | |
| Participation Coef., Modularity, Sex, Race, Ethnicity, Education, MeanFD, $MeanFD^2$, eTIV | -0.06 | 0.03 | -1.87 | 0.06 |
| **Accounting for graph theory metrics as moderators, and all other covariates** | | | | |
| Participation Coef.*age, Modularity*age, Participation Coef., Modularity, Sex, Race, Ethnicity, Education, MeanFD, $MeanFD^2$, eTIV | -0.05 | 0.06 | -0.92 | 0.36 |

Notes. All graph theory metrics were calculated using functional connectivity matrices generated from surface-based cortical data applied to the Schaefer atlas with volumetric subcortical data applied to the Cole atlas. Mean FD refers to the motion metric average framewise displacement. eTIV = estimated total intracranial volume.
